# Quantifying visual pathway overlap to identify training-related microstructural change in hemianopia

**DOI:** 10.64898/2026.07.28.740991

**Authors:** Miah Browne, Rebecca Millington-Truby, Holly Bridge, Arash Sahraie, Sara Ajina

## Abstract

**Background:** Visual training can improve residual vision in hemianopia, but there is significant variability in patient outcomes which may reflect differences in neural structures preserved after injury. Diffusion MRI tractography offers a means of assessing white-matter microstructure in vivo, however substantial overlap between neighbouring visual pathways complicates attribution of differences to specific tracts. We quantified tract overlap to identify relatively unique pathway segments to assess training-related microstructural change.

**Methods:** Diffusion MRI data was acquired in six participants with homonymous hemianopia before and after 3-6 months of visual training. Tractography was used to reconstruct three pathways implicated in residual vision: lateral geniculate nucleus (LGN)-V1, LGN-V5 and pulvinar-V5, together with secondary superior colliculus-LGN and superior colliculus-pulvinar pathways. Streamline overlap was quantified along neighbouring tracts, and comparison of pre- to post-training fractional anisotropy was restricted to unique or pair-specific non-overlapping segments. Exploratory relationships with visual outcomes were also examined.

**Results:** Fractional anisotropy increased after training in both early and late pair-specific segments of the LGN-V5 pathway in the lesioned hemisphere. In the late segment, this increase was significantly greater than in the corresponding unique LGN-V1 segment, whereas change in the early LGN-V5 segment did not differ significantly from the unique pulvinar-V5 segment. No significant training-related changes were observed in collicular pathways. Brain-behaviour analyses also preferentially implicated LGN-V5: baseline fractional anisotropy was strongly associated with post-training Gabor detection, and training-related fractional anisotropy change showed a positive association with improvement in Gabor detection, although the latter did not survive multiple comparison correction.

**Conclusion:** These findings support a role for the LGN-V5 pathway in visual plasticity after hemianopia and demonstrate the importance of explicitly quantifying tract overlap when attributing microstructural change to small, neighbouring white-matter pathways.

## INTRODUCTION

Homonymous hemianopia, or the loss of half of the visual field, is recognised as one of the most common functional deficits after a stroke, impacting 30-60% of stroke survivors (Rowe et al., 2013; Willis and Cavanaugh, 2023) and substantially affecting quality of life (Pollock et al., 2019; Rowe et al., 2022). In some patients, a degree of visual processing remains possible within the blind field, an ability sometimes referred to as ‘blindsight’ (Cowey, 2010; Weiskrantz et al., 1974). Residual visual function is typically strongest for high-contrast and moving stimuli (Das et al., 2014; Riddoch, 1917), although other preserved abilities including emotion processing or ‘affective blindsight’ (Celeghin et al., 2015; de Gelder et al., 1999) have also been described.

Understanding the visual features that most effectively engage residual processing has facilitated the development of visual training programs aimed at restoring a degree of function within the blind field, with increasingly promising results as rehabilitation strategies (reviewed in (Cavanaugh et al., 2025)). There is however still substantial variability in success between patients. While the reasons for this are not fully understood, this variability is likely to reflect heterogeneity in lesion location and extent, together with differences in the integrity of residual pathways and perilesional tissue available to support recovery (Barbot et al., 2021; Beh et al., 2022; Willis et al., 2025). Identifying the pathways that facilitate recovery is therefore crucial for guiding rehabilitation strategies.

Considerable uncertainty remains regarding the subcortical pathways that support residual visual processing after damage to the primary visual pathway. Proposed candidates include projections from the lateral geniculate nucleus (LGN) to the extrastriate motion area (V5) (Ajina et al., 2015; Ajina and Bridge, 2018; Bridge et al., 2010; Schmid et al., 2010; Willis et al., 2024), the pulvinar to V5 (Warner et al., 2012, 2010), and superior colliculus to extrastriate visual areas (Leh, 2006; Leh et al., 2006). Evidence has perhaps most consistently implicated the LGN-V5 pathway, although methodological challenges have limited the ability to investigate in detail the relative contributions of neighbouring tracts.

Diffusion MRI (dMRI) combined with tractography provides the principal non-invasive method for reconstructing white matter pathways in vivo and assessing their structural properties (Catani et al., 2012; Johansen-Berg, 2010; Jones et al., 2013). However, studies of residual vision have used a range of approaches to quantify pathway integrity. Early work focused on surrogate measures of ‘tract strength’ (Tamietto et al., 2012), and more recent studies have continued to report measures such as streamline count (Prabhakar et al., 2026), or fibre connectivity density (Sungkarat et al., 2024). Such measures offer some value in detecting gross degeneration or a wholly absent pathway (Sungkarat et al., 2024). However, they cannot be a surrogate for biological fibre density nor provide, in this context, meaningful inferences based on relative strength or weakness. Streamline counts are influenced by multiple factors unrelated to the underlying tissue architecture, including seed and target size or placement, tract length and curvature, crossing fibres, tracking and stopping criteria, lesion-related uncertainty in fibre orientation estimates, and the number of samples generated. Indeed, these limitations have motivated development of quantitative tractography approaches (Smith et al., 2015, 2013). To derive more biological meaningful differences in pathway density, one needs to apply dMRI signal-informed filtering or fixel-based approaches (Raffelt et al., 2015). Unfortunately, this can require scan acquisitions with higher b-values and, in some cases, multi-shell data that is not always available in clinical cohorts.

An alternative approach is to assess diffusion properties within reconstructed white-matter bundles (Ajina et al., 2015; Sampaio-Baptista et al., 2013). Fractional anisotropy (FA) is a sensitive but non-specific marker of white-matter microstructure, influenced by factors including axonal packing and density, myelination, axonal calibre, fibre coherence and orientation dispersion (Henriques et al., 2023). Although FA cannot distinguish between these underlying biological processes, tract-specific longitudinal measurements can provide a sensitive index of microstructural change, particularly when comparisons are made within individuals and spatial correspondence and tract definition are carefully controlled (Sampaio-Baptista et al., 2013).

Tract-profile and segment-wise analyses have further enabled microstructural measurements to be localised to particular portions of a pathway rather than averaged across the tract as a whole (Ajina et al., 2015; Willis et al., 2024). This is especially important when comparing where neighbouring pathways overlap. However, the challenge of dissociating the relative contributions of overlapping visual pathways remains. Previous studies attempting to separate LGN-V1 from LGN-V5 pathways selected tract segments without explicitly quantifying the degree of streamline overlap (Ajina et al., 2015; Willis et al., 2024). Furthermore, studies of residual vision and visual training in hemianopia have focussed on comparisons of LGN-V5 and LGN-V1 pathways (Willis et al., 2024). To our knowledge, no previous study has directly compared training-related dMRI changes in LGN-V5 and pulvinar-V5 pathways in this population. This is primarily because LGN and pulvinar projections to V5 are known to show substantial spatial overlap along their trajectory (Allen et al., 2015; Rowe et al., 2023).

The present study addresses this problem by explicitly quantifying overlap between the candidate visual pathways before assessing training-related microstructural change. dMRI data was acquired in six participants with homonymous hemianopia before and after completing 3-6 months of visual training, as previously described (Ajina et al., 2021). Tractography was used to reconstruct three pathways implicated in residual vision: LGN-V1, LGN-V5, and pulvinar-V5. We additionally reconstructed two pathways connecting the superior colliculus with LGN and pulvinar, since any contribution of retinal input to V5 via the superior colliculus is likely to involve at least one intervening subcortical relay (Berman and Wurtz, 2008; Torrealba et al., 1981). Critically, streamline overlap was quantified between neighbouring tracts, and microstructural analyses were restricted to pathway segments identified as relatively unique. This approach allowed us to test whether specific visual pathways showed training-related microstructural change, and whether the changes were associated with improvements in visual function, while explicitly accounting for anatomical overlap between tracts.

## MATERIALS AND METHODS

### Participants

The participants in this study are described in a previous publication (Ajina et al., 2021). To summarise, six participants with homonymous hemianopia or quadrantanopia (3 female) took part in the training study. All had sustained a stroke at least 6 months before enrolling in the study. Average age at the time of participation was 61.3 ± 13.1 years and average time since lesion at first scan was 14 ± 7 months. Written, informed consent was obtained from all participants and the research adhered to the Declaration of Helsinki. Ethical approval was provided by the Oxfordshire Research Ethics Committee (Reference B 08/H0605/156) or Oxford University Central Ethics Committee (Reference MS-IDREC-C2-2015-025).

### Study design

Training participants attended two study sessions, before and after a period of visual training lasting several months. Each session included a 3T MRI scan, visual field testing and visual psychophysics. After the first session, participants were provided with training apparatus in their home and undertook a visual training paradigm provided by Neuro-Eye Therapy (see (Sahraie et al., 2013) for details). After a period of 4-8 months, participants returned to University of Oxford to undergo the second session (average time post-pre: 6.8 months, ± 3.6 months S.D.).

### Training protocol

The training procedure was conducted in the participant’s home on an IBM compatible personal computer, mounted on a frame (Figure 1A). Gamma corrections were conducted on all monitors using an LMT luminance meter (LS100; Konica Minolta, Inc, Tokyo) at 256 equistepped logical colours. Participants sat with a chin rest 40 cm from the monitor, with line of sight approximately level with the fixation point. Viewing was binocular throughout the experiment.

**Figure 1.**
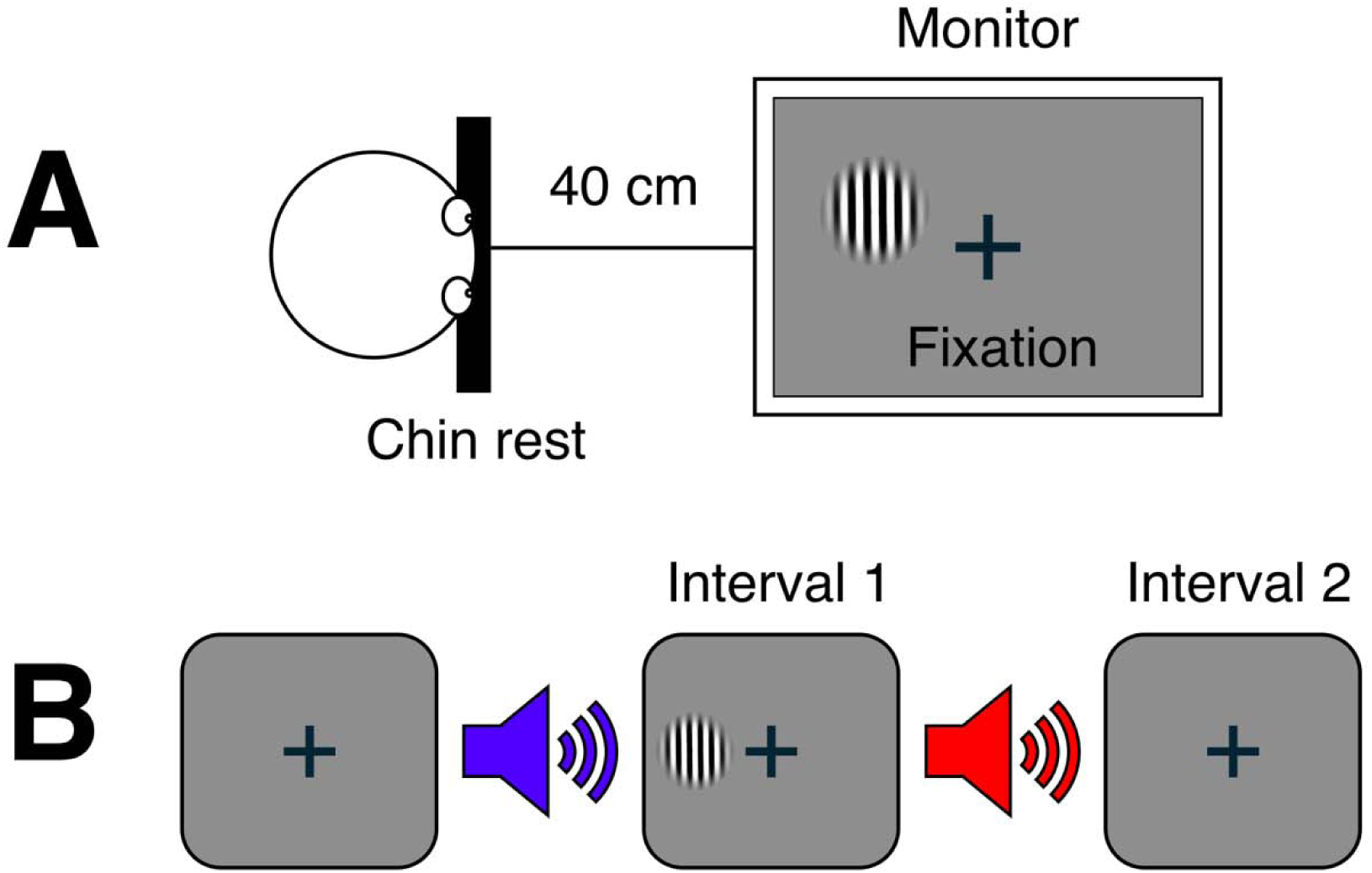
Visual training apparatus and procedure. Participants sat with their head on a chin rest at a distance of 40 cm from the monitor, mounted on a frame. **B.** The training task consisted of a 2-AFC temporal detection task in which a spatially and temporally modulated Gabor patch was presented in one of two intervals while the participant maintained central fixation. The target contrast was algorithmically controlled to maintain difficulty. Positive auditory feedback was provided if they were correct. Each training session lasted 25 min, with a 3 min rest halfway through the session.

Training stimuli consisted of vertically-oriented achromatic Gabor patches, with spatial smoothing of the boundaries (spatial frequency=1cycle/°; temporal frequency=10 Hz; diameter=6°). Stimuli were presented at three predetermined retinal eccentricities in a randomly interleaved order, with the exact locations tailored to each participant’s deficit (see (Ajina et al., 2021) Table 1 for coordinates). One of the three locations overlapped with the test stimulus in psychophysical experiments. This represented the primary target of the visual training study and was the location for all psychophysical assessments, performed before and after the training programme.

**Table 1.**
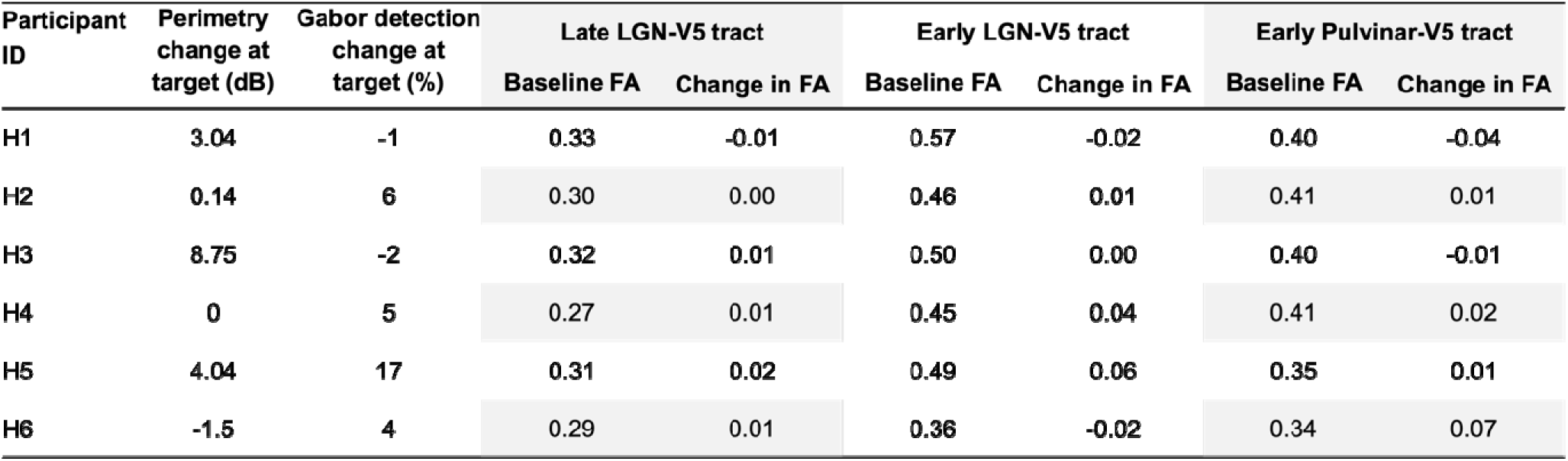
Individual visual outcomes and fractional anisotropy (FA) measures in pathway segments used for brain-behaviour analyses. Change in clinical perimetry represents the pre- to post-training change in visual field sensitivity within the trained region, expressed in decibels (dB). Change in Gabor detection represents the pre- to post-training change in detection performance at the trained target location. Baseline FA and change in FA (post-training minus pre-training) are shown for the late and early segments of the LGN-V5 pathway and the early segment of the pulvinar-V5 pathway in the lesioned hemisphere. Tract segments were selected according to the overlap criteria described in the Methods.

| Participant ID | Perimetry change at target (dB) | Gabor detection change at target (%) | Late LGN-V5 tract |  | Early LGN-V5 tract |  | Early Pulvinar-V5 tract |  |
| --- | --- | --- | --- | --- | --- | --- | --- | --- |
|  |  |  | Baseline FA | Change in FA | Baseline FA | Change in FA | Baseline FA | Change in FA |
| H1 | 3.04 | -1 | 0.33 | -0.01 | 0.57 | -0.02 | 0.40 | -0.04 |
| H2 | 0.14 | 6 | 0.30 | 0.00 | 0.46 | 0.01 | 0.41 | 0.01 |
| H3 | 8.75 | -2 | 0.32 | 0.01 | 0.50 | 0.00 | 0.40 | -0.01 |
| H4 | 0 | 5 | 0.27 | 0.01 | 0.45 | 0.04 | 0.41 | 0.02 |
| H5 | 4.04 | 17 | 0.31 | 0.02 | 0.49 | 0.06 | 0.35 | 0.01 |
| H6 | -1.5 | 4 | 0.29 | 0.01 | 0.36 | -0.02 | 0.34 | 0.07 |

The training task required detection of a Gabor patch using a temporal two-alternative forced-choice (2-AFC) task. Participants were required to report whether a target stimulus was presented during the first or second of two intervals. The intervals were separated by auditory cues of “one” or “two”, announcing the start of each interval. Each trial lasted 6 s in total and finished with a low tone (Figure 1B). At the start of training the target contrast at all three locations were set to 95%. This contrast at each location was then lowered by 10% after three consecutive sessions when correct performance was above 84%. Reduction of performance to 64% and below resulted in an increase of contrast by 5% in the subsequent training session. This method has been shown to ensure maximum stimulation, while increasing task difficulty with improved performance (Sahraie et al., 2010). Auditory feedback was provided to denote correct discrimination.

### Psychophysical testing

This was a lab-based assessment, carried out during the two study sessions before and after the training period. Gabor detection was taken as an average score for detection of a drifting achromatic Gabor patch in a two-alternate forced choice protocol (temporal frequency 10Hz, spatial frequency 1.3 cycles/deg) across a range of luminance contrast levels (1%, 5%, 10%, 50%, 100%). This resulted in two mean scores per participant, (1) Gabor detection pre-training and (2) Gabor detection post-training. These scores were used to generate a single score representing change in Gabor detection with training (Table 1). For additional details, please see (Ajina et al., 2021).

To summarise the results, 2/6 participants could already perform Gabor detection at maximum contrast levels at baseline (prior to training) and thus showed little change with training (H1 and H3). All other participants showed variable improvement with training (Table 1).

### Visual Field testing

Static visual fields in the central 30° were measured before and after training in all participants using Humphrey Visual Field Analyser. We used the mean change in sensitivity in the trained region of the visual field, determined previously for each participant (see (Ajina et al., 2021) Table 2). According to this metric, 4/6 participants showed an increase in sensitivity after training and 2/6 participants showed either no change or a deterioration (Table 1). There was a significant effect of training across the group (mean gain of 3.4dB, one-tailed Wilcoxen signed rank test; sum of ranks=17; no. pairs=7; p=0.047), which was absent in an equivalent untrained region of the affected hemifield.

### Magnetic Resonance Imaging

Participants were scanned either on a 3T Siemens Verio scanner or a 3T Siemens Prisma, using a 32-channel head coil, with the same scanner used for both visits. A T1-weighted 1mm^3^ isotropic resolution MPRAGE anatomical scan (TE=4.68ms, TR=2040ms; field of view=200 mm, flip angle=8deg) was acquired for each participant at each scan session.

Diffusion-weighted data were acquired using echo planar imaging (EPI; TR = 8900 ms, TE = 91.2 ms, and voxel size of 2×2×2 mm^3^). The diffusion weighting was isotropically distributed along 60 directions (b-value = 1500 s/mm^2^), and a non-DWI (B0) image was acquired every 16 volumes (total of four B0 volumes per image set). To minimise geometric distortions, two sets of images were acquired with the phase-encoded direction reversed. Each image set was corrected for motion and eddy-current related distortions, before running topup to correct for geometric distortions. Corrections were performed using tools from FSL (FMRIB Centre Software Library, Oxford University; http://www.fmrib.ox.ac.uk/fsl/), with subsequent steps in tensor modelling and tractography using MRtrix3 v3.0 (Tournier et al., 2019) diffusion MRI software suite. The T1-weighted structural scan was aligned to dMRI images using FLIRT and FNIRT.

### Region of interest definition

Cortical parcellation of V5 and V1 were performed using the Human Connectome Project multimodal parcellation (HCP-MMP) atlas (Glasser et al., 2016). Subcortical regions LGN and inferior pulvinar were obtained from a standard space atlas based on the Bayesian segmentation algorithm implemented in FreeSurfer (Iglesias et al., 2018). Superior colliculus masks were drawn manually on individual T1 scans.

### Streamline generation

Tissue response functions were estimated using the unsupervised ‘dhollander’ algorithm (Dhollander et al., 2016; Tournier et al., 2019). Single-shell 3-tissue constrained spherical deconvolution was then performed using MRtrix3Tissue to estimate fibre orientation distribution (FOD) of white matter-like, grey-matter-like, and CSF-like tissue compartments (Dhollander and Connelly, 2016). The resulting tissue-component images were intensity-normalised to correct intensity inhomogeneities. Streamlines were then generated using tckgen with the following parameters: iFOD2 algorithm, 3000 streamlines, with default settings for minimum and maximum length, step size, and maximum angle (Smith et al., 2012; Tournier et al., 2010). Tracts were cleaned to remove outlier streamlines by removing any streamlines greater than two standard deviations above mean length for each tract using tckedit. Tracts were then resampled to 100 nodes distributed equally along the length of the tract (Yeatman et al., 2012) using tckresample. This resampling allowed us to perform standardised comparisons across participants and tracts.

### Fractional Anisotropy

Although the diffusion tensor model (Basser et al., 1994; Pierpaoli and Basser, 1996) can be inappropriate for tracking, it can provide an accurate representation of the signal and its statistics (Rokem et al., 2015). The tensor model was fitted at each voxel using dwi2tensor to derive FA maps, from which the mean and variation along any fascicle bundle could be calculated. FA provides a measure of the directionality of water molecule movement, which relates to the geometric organization of axons and fascicles in each voxel (e.g. crossing, merging or ‘kissing’ fibres), the degree of myelination of axons in the white matter (Beaulieu and Allen, 1994), and their packing density (Sen and Basser, 2005). In cases of brain damage, a decrease in FA can be indicative of loss of structural integrity of fibre bundles (Jones et al., 2013), such as Wallerian degeneration (Beaulieu et al., 1996). Conversely, increases in FA can be an indication of plasticity in white matter tracts due to increased myelination and fibre packing density (Sampaio-Baptista et al., 2013; Sen and Basser, 2005).

### Overlap of adjoining pathways

Dice coefficients were calculated to estimate the overlap of the cleaned, resampled tracts. This was done by generating streamline count maps to represent the different tracts in native space using tckmap. Two approaches were used: (1) Weighting by streamline density to give a weighted dice score that is equivalent to tract FA measurements averaged across streamlines (2) A non-weighted voxel position mask reflective of *all* streamlines as a binary template to determine the size of the intersection relative to total extent. Although arguably this would be less meaningful for FA metrics, the latter method provides less conservative estimates. With both methods, an average was taken across pre and post-training scans.

There is no universally accepted estimate for dice coefficient when comparing adjoining but distinct neural pathways. However, dice scores revealed significant overlap between LGN-V5 and Pulvinar-V5 tracts (Mean Dice 0.7 ± SD 0.1 using more conservative weighted method) as well as between LGN-V1 and LGN-V5 tracts (Mean Dice 0.4 ± SD 0.1). Of note, both approaches gave similar results, suggesting binary masks would be suitable as a proxy for streamline trajectory.

Further analysis looked for specific regions along the three pathways that might demonstrate anatomical specificity. To do this, we created a pipeline to estimate the proportion of streamlines in each tract that overlapped with the other two pathways along its trajectory from subcortical to cortical endpoints. We first ensured that all tract bundles were aligned in a consistent direction from subcortical to cortical ROI using custom built scripts. We then used the binary template for each tract in native space to test for an overlap with streamlines in each tract of interest. Each individual streamline was tested at every point along its 100-node trajectory to quantify its overlap with either of the other two pathways. This was used to calculate the percentage of streamlines at each node that were either (1) ‘unique’ i.e. did not overlap with either other pathway, (2) overlapped with one pathway but not the other, (3) overlapped with the other pathway but not the first, (4) overlapped with both pathways.

Plots demonstrating tract overlap (Figure 2) were then used to define tract segments for fractional anisotropy (FA) measures. We applied a conservative threshold of at least 95% unique streamlines, meaning that fewer than 5% of streamlines overlapped with either of the other tracts for the region to be classified as unique. Using this criterion, nodes 1-5 were selected to represent the unique portion of the pulvinar-V5 tract, and nodes 88-100 were selected to represent the unique portion of LGN-V1.

**Figure 2.**
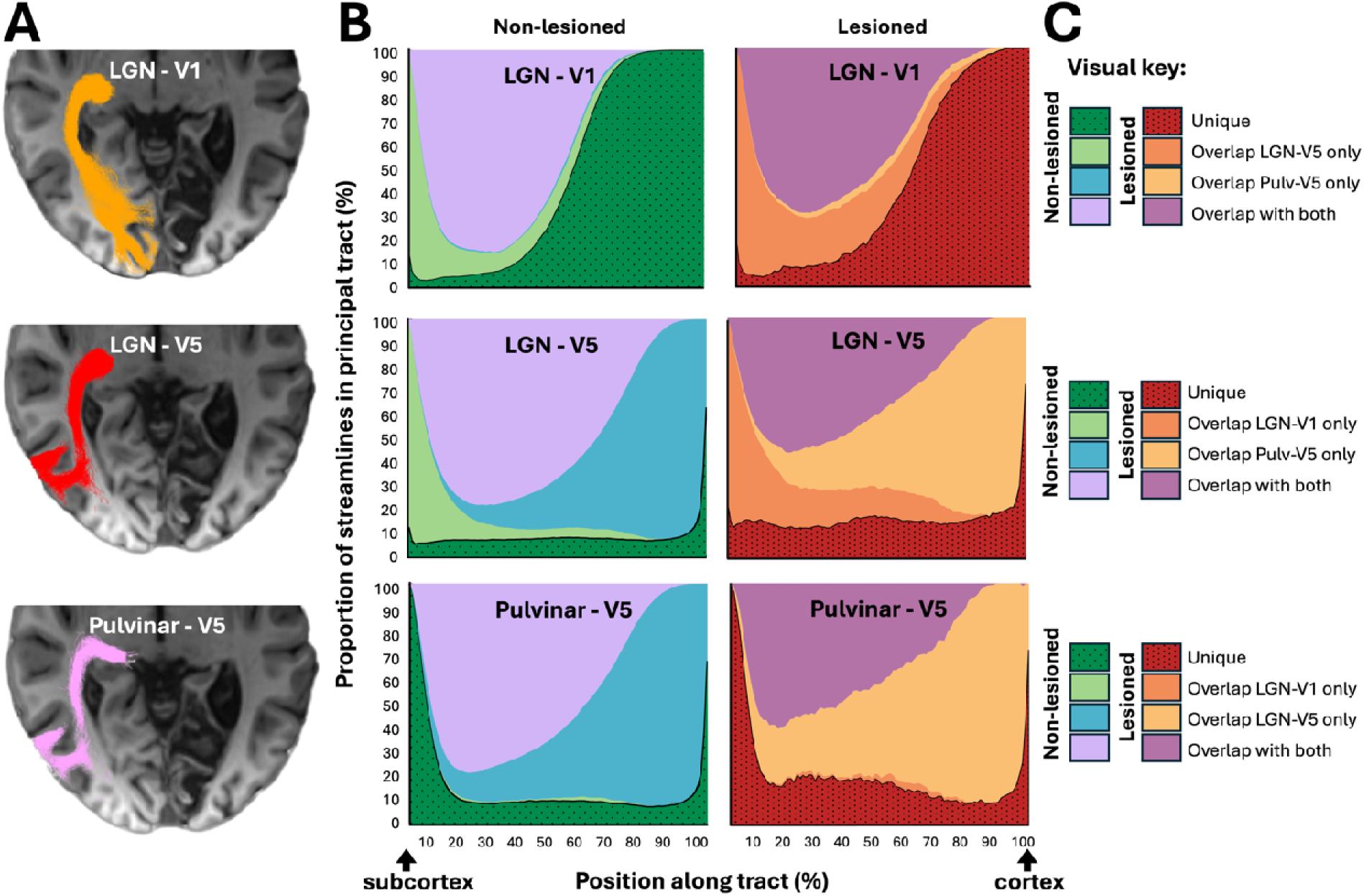
Overlap of the three principal visual tracts involving optic radiation. **(A)** Representative reconstruction of LGN-V1, LGN-V5 and pulvinar-V5 pathways in participant H4. **(B)** For each pathway separately, the proportion of streamlines at each node along the tract that did not overlap with either neighbouring pathway, overlapped with one neighbouring pathway only, or overlapped with both neighbouring pathways. Position along each tract was normalised from 1 (subcortex) to 100% (cortex), and results are shown separately for the lesioned and non-lesioned hemispheres. **(C)** Visual key illustrating, for each principal tract, the classification of streamline overlap categories used in **B**, including streamlines unique to that pathway, overlapping with either neighbouring pathway alone, or overlapping with both neighbouring pathways.

No segment of the LGN-V5 tract achieved 95% uniqueness relative to both neighbouring tracts simultaneously. However, the same threshold could be used in a pair-specific manner to identify LGN-V5 segments with selective non-overlap. Specifically, nodes 1-5 were selected to represent the early LGN-V5 segment that showed at least 95% non-overlap with pulvinar-V5, but substantial overlap with LGN-V1, while nodes 88-100 were selected to represent the late LGN-V5 segment that showed at least 95% non-overlap with LGN-V1, despite substantial overlap with pulvinar-V5. This allowed direct comparison of these pair-specific LGN-V5 segments with the fully unique segments of pulvinar-V5 and LGN-V1, respectively, to assess the specificity of microstructural changes.

In addition to the three principal tracts of interest, we carried out secondary analyses in subcortical pathways between (i) superior colliculus and LGN and (ii) superior colliculus and pulvinar. Involvement of superior colliculus in blindsight is well documented, and it is reasonable to suppose that superior colliculus and its connections may be involved in visual training tasks targeting blindsight. Collicular tracts projecting directly to extrastriate cortex have been shown to be unreliable in diffusion tracking software (Ajina et al., 2015; Sangchooli et al., 2026). This is likely due to limitations when passing through grey matter way-points reflecting disynaptic pathways (Berman and Wurtz, 2010). As an alternative approach, tracts appear more robust and reproducible by running streamlines separately from superior colliculus to LGN or pulvinar before then tracking to visual cortex, thus representing two components of a probable di-synaptic pathway. Since both tracts share an end-point, they will naturally overlap towards that terminus. Thus to account for this, a similar approach was taken as for principal tracts to quantify the overlap along their trajectory in order to identify specific regions. As subcortical pathways were considerably shorter than pathways between subcortex and visual cortex, a less conservative threshold of >=80% streamlines was applied resulting in nodes 70-100 being used to represent SC-LGN and SC-pulvinar tracts (see supplemental Figure 1).

### Statistics / Tract-based statistics

Groupwise analyses tested for training-related change in fractional anisotropy (FA) across the three principal and two secondary tracts. To adopt an anatomically driven approach that accounted for overlap between pathways, primary analyses were restricted to the tract segments identified from Figure 2 using the overlap criteria described above. Paired two-tailed t-tests were used to test for change in FA between pre- and post-training. Because no segment of LGN-V5 met the criterion for uniqueness relative to both neighbouring pathways, additional pair-specific analyses were performed for LGN-V5. These compared FA change in the early LGN-V5 segment, which showed minimal overlap with pulvinar-V5, against the unique pulvinar-V5 segment, and FA change in the late LGN-V5 segment, which showed minimal overlap with LGN- V1, against the unique LGN-V1 segment. This allowed tract-specific comparison of LGN-V5 with each overlapping pathway while retaining the same conservative overlap threshold. Equivalent analyses were performed for the contralesional hemisphere.

We were also interested in whether visual improvement post-training was related to either visual pathway microstructure at baseline or change in FA post-pre training. Correlation analyses used the same mean FA that was used to test for an effect of training above. FA was averaged along unique segments for each visual tract to generate tract-specific scores for pulvinar-V5 and LGN- V1 tracts in each participant. For the LGN-V5 tract, this was limited since the early LGN-V5 segment was unique from pulvinar-V5 but not LGN-V1, while the late LGN-V5 segment was unique from LGN-V1 but not pulvinar-V5. Primary correlation analyses focused on change in clinical Perimetry score and change in Gabor detection as markers of positive visual outcomes. However, we also looked at post-training Gabor detection, to account for patients who were at ceiling on this measure pre-training. For all correlation analyses, FDR correction was applied.

## Results

### 1. FA in ipsilesional LGN-V5 tract increases after visual training

In the lesioned hemisphere (Figure 3), both early and late pair-specific LGN-V5 segments showed a significant increase in fractional anisotropy following training (early paired t = 2.7, p = 0.01, df = 29; late paired t = 3.2, p = 0.002, df = 77). To assess for tract specificity, change in FA within these LGN-V5 segments was compared directly with change in FA in the corresponding non-overlapping tract region. For the late segment, FA change in LGN-V5 differed significantly from the unique LGN-V1 segment (t = 3.45, p = 0.0009, df = 77), with mean FA increasing in LGN-V5 but decreasing in LGN-V1 after training. In contrast, change in FA in the early LGN-V5 segment showed no significant difference when compared to the unique pulvinar-V5 segment (t = 1.1, p = 0.28, df = 29).

**Figure 3.**
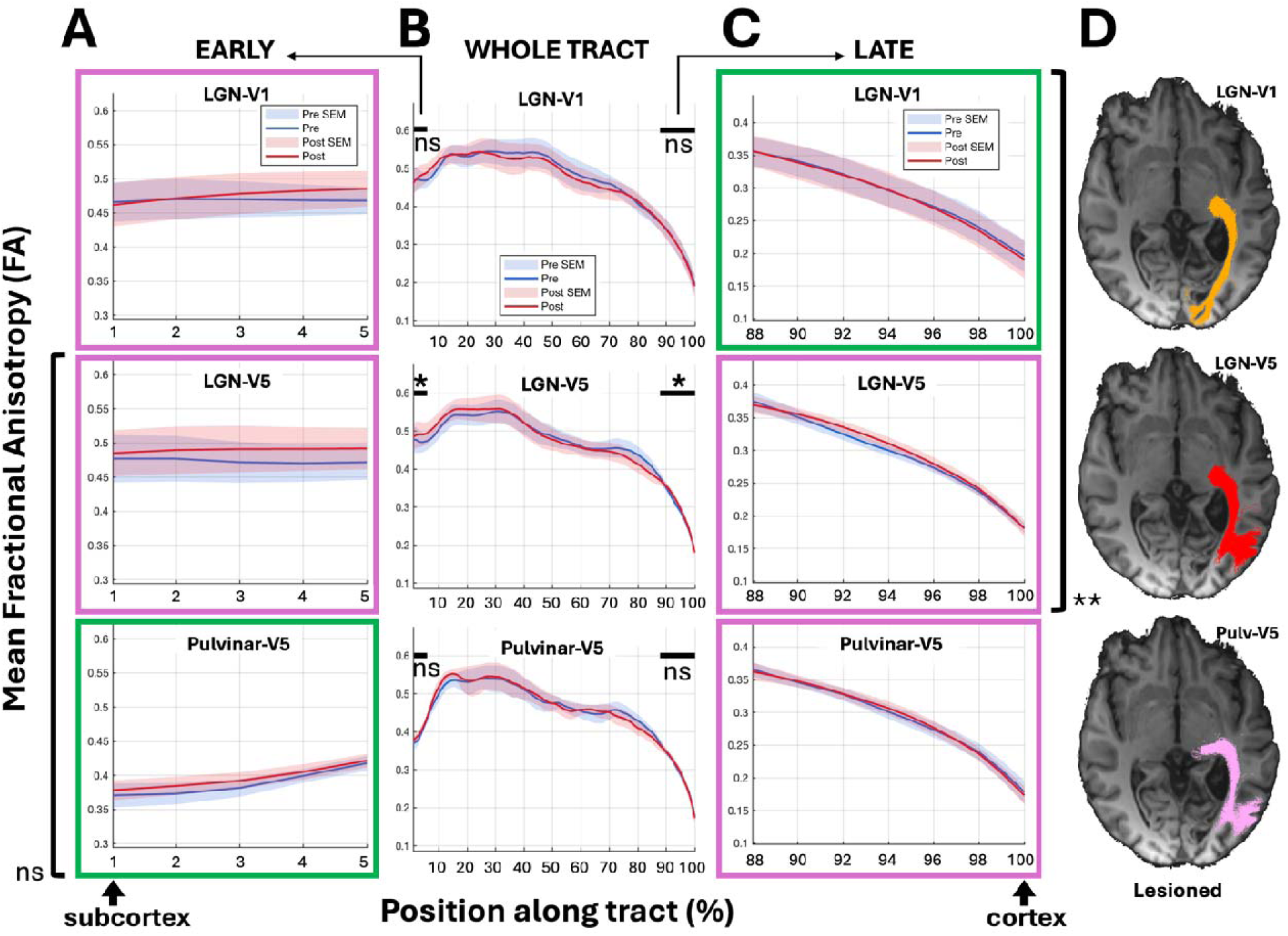
Fractional anisotropy (FA) along the principal visual tracts in the lesioned hemisphere before and after visual training, from subcortex (1%) to cortex (100%) endpoints. **A**. Early component shows the first 5% of tracts, closest to the subcortical origin. Pre-training FA is plotted in blue, post-training in red. Standard error of the mean shown by shaded regions. Unique tract segments with > 95% non-overlapping streamlines compared to *both* other pathways are highlighted in a green box, all other segments are in a pink box. Since LGN-V5 tracts do not show any ‘unique’ regions along its tract, pairwise comparisons to non-overlapping tracts are depicted with black vertical lines. **B.** Whole tract profiles. Regions selected for expansion in panels A (early) and C (late) are indicated by black horizontal bars above the plots. Tract segments with significant FA change post-pre training are indicated with an Asterix. **C.** Late component shows the final 12% of tract profiles, where LGN-V1 streamlines (top row) are >95% unique from LGN-V5 and Pulvinar-V5 streamlines. ns = non-significant, * p 0.01, ** p < 0.001. **D.** Representative reconstruction of pathways in participant H4 overlaid on co-registered T1w scan.

In the non-lesioned hemisphere (Figure 4), all tracts showed a small decrease in FA after visual training. Only the LGN-V1 tract reached significance (early paired t = 2.9, p = 0.006, df = 34, late paired t = 2.3, p = 0.02, df = 90). However, the change in FA was not greater than seen in LGN-V5 tract (paired t = 1.0, p = 0.34, df = 90), suggesting the effect was relatively weak and, to some extent, was ubiquitous in other pathways. For analyses of subcortical pathways connecting superior colliculus to pulvinar or LGN, there were no significant changes to FA with training, in either the lesioned or non-lesioned hemisphere of both pathways (Supplementary Figure 2).

**Figure 4.**
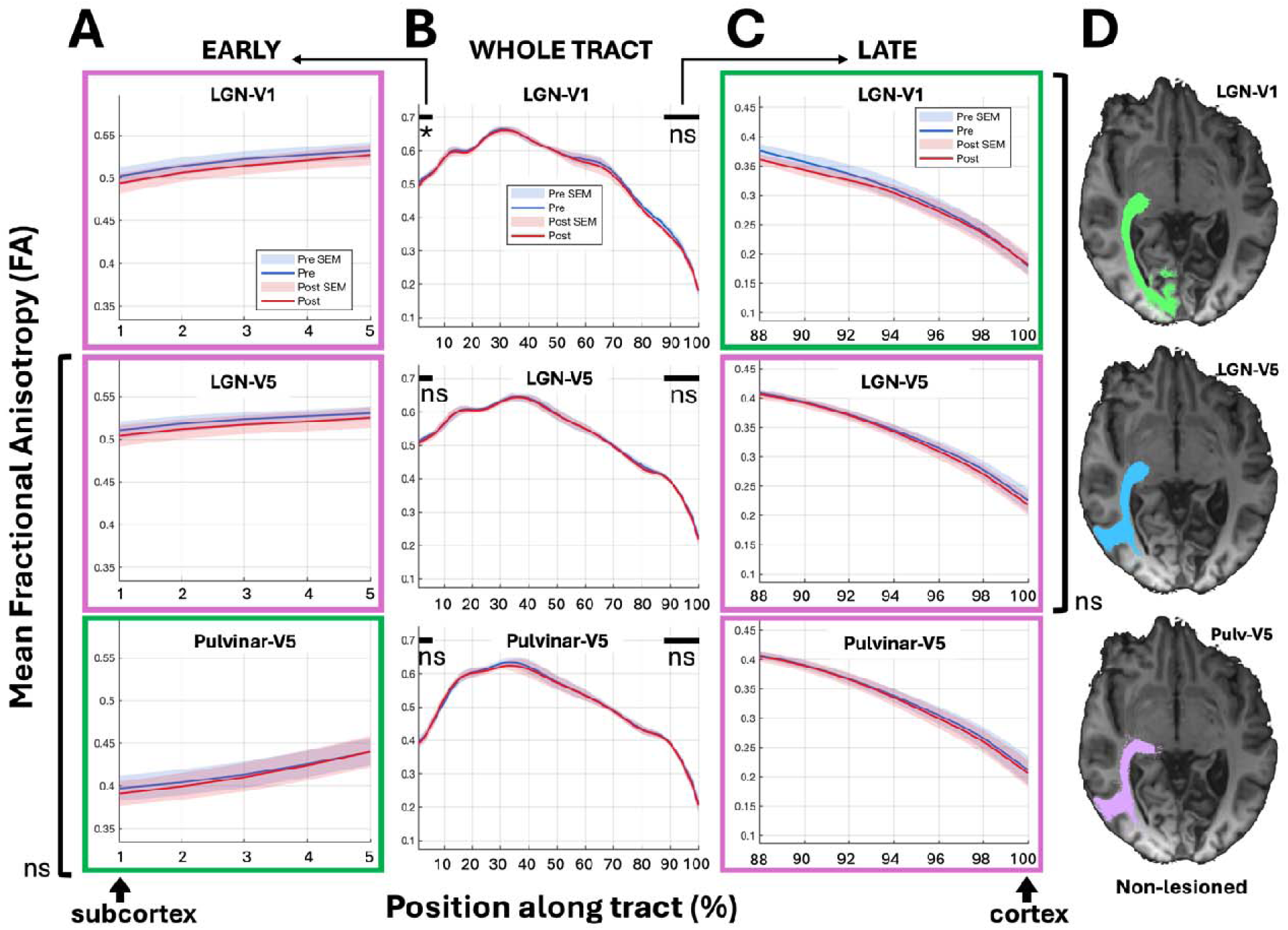
Fractional anisotropy (FA) along the principal visual tracts in the non-lesioned hemisphere before and after visual training, from subcortex (1%) to cortex (100%) endpoints. **A**. Early component shows the first 5% of tracts, closest to the subcortical origin. Pre-training FA is plotted in blue, post-training in red. Standard error of the mean shown by shaded regions. Unique tract segments with > 95% non-overlapping streamlines compared to *both* other pathways are highlighted in a green box, all other segments are in a pink box. Since LGN-V5 tracts do not show any ‘unique’ regions along its tract, pairwise comparisons to non-overlapping tracts are depicted with black vertical lines. **B.** Whole tract profiles. Regions selected for expansion in panels A (early) and C (late) are indicated by black horizontal bars above the plots. Tract segments with significant FA change post-pre training are indicated with an Asterix. **C.** Late component shows the final 12% of tract profiles, where LGN-V1 streamlines (top row) are >95% unique from LGN-V5 and Pulvinar-V5 streamlines. ns = non-significant, * p 0.01. **D.** Representative reconstruction of pathways in participant H4 overlaid on co-registered T1w scan.

### 2. LGN-V5 tract integrity is associated with visual outcome after training

Although the sample size in this study was small, exploratory analyses examined whether (1) baseline microstructure pre-training might predict visual improvement, and (2) training-related visual improvement was associated with a change in microstructure, namely an increase in FA. Visual outcomes for correlation analyses included change in clinical Perimetry score and change in Gabor detection, as well as post-training Gabor detection to account for patients at ceiling pre-training.

Brain-behaviour analyses for clinical Perimetry (Figure 5, middle column) revealed notable but non-significant positive associations with baseline FA in the pair-specific LGN-V5 segments, with the strongest association observed in the late segment (r = 0.77, Figure 5B and C). Conversely, baseline FA was only weakly associated with change in Gabor detection across all examined pathways (mean r = -0.14 ± 0.16 SE). There was, however, a strong association between baseline FA and post-training Gabor detection in the early pair-specific LGN-V5 segment (r = 0.96, FDR p = 0.016). A positive association was also demonstrable in the late pair-specific LGN-V5 segment, although this did not reach statistical significance (r = 0.78). None of the other tracts at baseline showed a significant association with visual outcome (mean r = 0.3, ± 0.13 SE).

**Figure 5.**
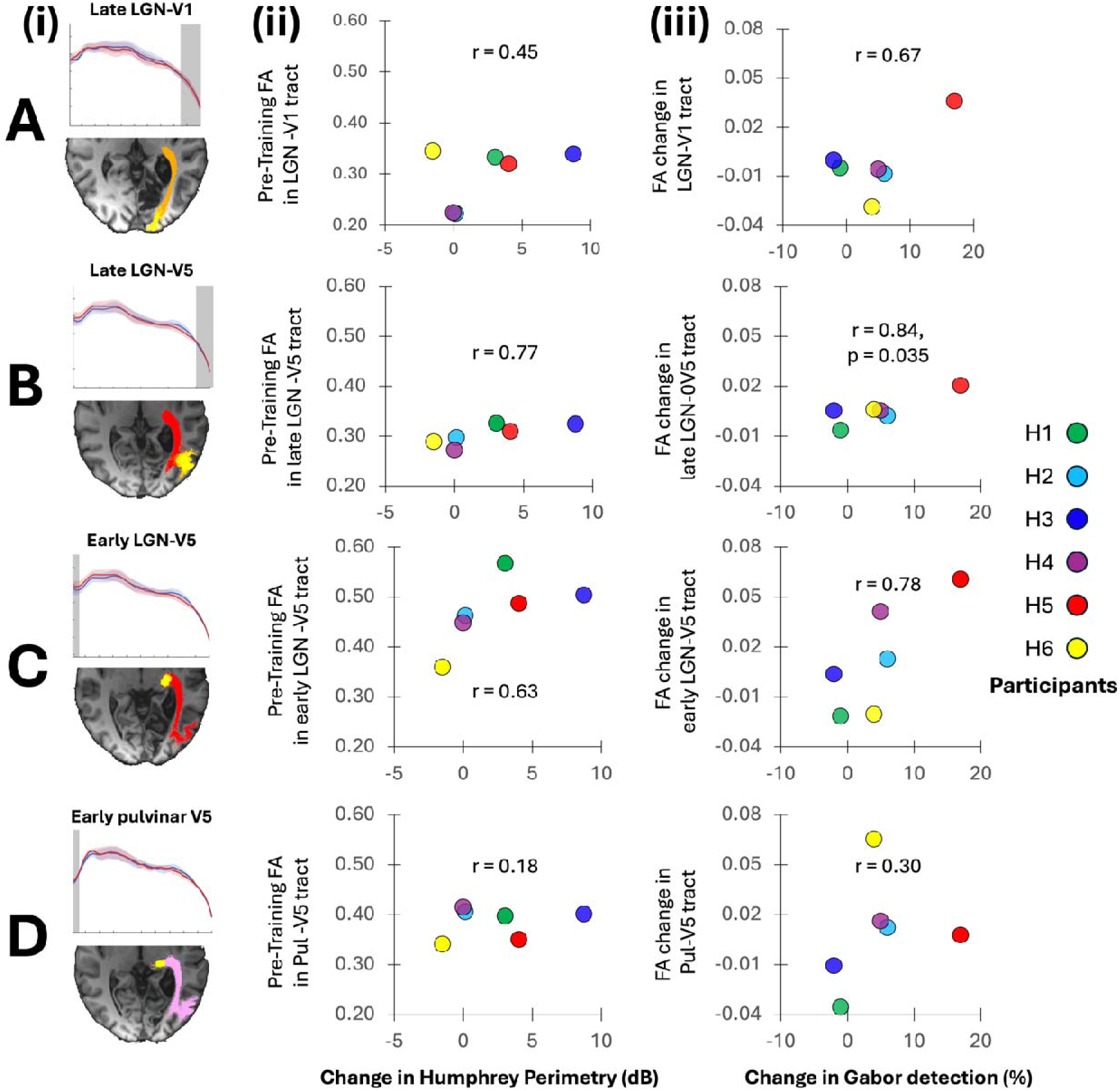
Brain-behaviour correlations in ipsilesional visual tracts. (i) Left column: Visualisation of tract profiles and representative streamlines in H4, highlighting the ‘unique’ or pair-specific portion of each tract used for correlation analyses in yellow. **(ii)** Middle column: Correlation between change in Humphrey Perimetry in the trained region of the visual field versus FA at baseline. (iii) Right column: Correlation between change in Gabor detection (%) versus FA change. **A.** Late LGN-V1 tract, **B.** Late LGN-V5 tract, **C.** Early LGN-V5 tract, **D.** Early pulvinar-V5 tract. Individual participant colour codes shown to the right. P values provided where p < 0.05.

For the analysis of FA change following visual training, we found that improvement in the Gabor detection task (Figure 5, right column) was positively associated with FA change in the LGN-V5 tract in the damaged hemisphere, in particular the late portion (r = 0.84, Fig 5C), although the effect did not survive correction for multiple comparisons. An equivalent association was not observed with the other tracts. Similarly, none of the tracts showed a significant association between change in Perimetry and change in FA, with the strongest association seen in the LGN- V1 tract (r = 0.51, mean r = 0.00 ± 0.19 SE across all other tracts).

## Discussion

This study provides, to our knowledge, the first tract-specific analysis of training-related microstructural change in posterior visual pathways in hemianopia that explicitly quantifies and accounts for overlap between neighbouring tractography reconstructions. This enabled us to compare three closely adjacent visual pathways: LGN-V1, LGN-V5 and pulvinar-V5. By measuring streamline overlap along each tract and restricting analyses to pathway segments with minimal overlap, we found that increases in fractional anisotropy following visual training were most evident in the LGN-V5 pathway of the lesioned hemisphere. Brain-behaviour relationships also preferentially implicated this pathway, with baseline FA and FA change in LGN-V5 showing the strongest associations with visual outcome measures. Together, these findings support a role for geniculo-extrastriate circuitry in visual recovery training after stroke, while also highlighting the importance of anatomically informed analyses when neighbouring visual pathways cannot be cleanly separated using conventional diffusion tractography.

## Quantifying overlap to isolate pathway-specific regions

A central aim of the present study was to address tract overlap explicitly. Small visual pathways within the optic radiation are difficult to dissociate using diffusion MRI because they travel in close proximity and may share voxels across substantial portions of their trajectory (for review, see (Ajina et al., n.d.). If microstructural estimates are averaged across an entire reconstructed tract, pathway-specific effects may be diluted or misattributed to neighbouring bundles. Although this issue has been recognised previously, prior studies have generally inferred rather than quantified its extent or have tried to take steps to avoid it without necessarily estimating it (Rowe et al., 2023; Willis et al., 2024).

Here, we found that geniculate and pulvinar projections to V5 showed considerable overlap, particularly towards their shared cortical endpoint. This makes it difficult to attribute whole-tract microstructural metrics to either pathway with confidence. In this context, examining tract segments further from the shared cortical target, where overlap is reduced, provides a more conservative approach. Using this strategy, we identified portions of pulvinar-V5 and LGN-V1 that were largely unique, as well as pair-specific segments of LGN-V5 that showed minimal overlap with one neighbouring pathway, although no LGN-V5 segment was fully unique relative to both neighbouring tracts simultaneously.

This finding is important because it illustrates both the value and the limitation of tract-specific analyses in this region. On the one hand, quantifying overlap allowed us to test whether training-related FA changes were preferentially associated with LGN-V5 rather than adjacent pathways. On the other hand, the absence of a fully unique LGN-V5 segment means that the results should not be interpreted as definitive evidence for a completely isolated anatomical pathway. Rather, they suggest that the strongest training-related microstructural change occurs within the reconstructed geniculo-extrastriate trajectory, even after accounting for its overlap with neighbouring LGN-V1 and pulvinar-V5 pathways.

Broader anatomical connectivity further complicates interpretation. V5 has widespread reciprocal connections with visual cortex, including V1-V4, MST and VIP, and is linked to frontal visuomotor circuitry through connections between extrastriate visual areas and the frontal eye fields (Maunsell and van Essen, 1983; Schall et al., 1995). Emerging evidence also suggests that MT/V5-related pathways may interact with broader frontoparietal networks involved in attention and visuomotor control (Millington-Truby et al., 2026). Consequently, microstructural measures from any single reconstructed bundle should not be assumed to reflect one monosynaptic anatomical pathway in isolation. Tractogram-filtering and forward-modelling approaches, including SIFT (Smith et al., 2013), SIFT2 (Smith et al., 2015) and LiFE (Pestilli et al., 2014), can improve the biological plausibility or quantitative interpretation of tractography by relating streamline reconstructions to the diffusion signal or underlying fibre-density estimates.

However, these methods remain dependent on the candidate tractogram, and whole-brain tractograms may provide too few streamlines for reliable quantification of small or low-probability pathways, necessitating local ROI to ROI tractography (Rowe et al., 2023).

## LGN-V5 changes with visual training

The clearest training-related effect was an increase in FA within the LGN-V5 pathway of the lesioned hemisphere. This was observed in both early and late pair-specific LGN-V5 segments, suggesting that the finding was not restricted to a single portion of the reconstruction. Importantly, the late LGN-V5 segment showed a significantly greater FA increase than the corresponding unique LGN-V1 segment, supporting some degree of pathway specificity. The comparison with pulvinar-V5 was less clear, as FA change in the early LGN-V5 segment did not differ significantly from the unique pulvinar-V5 segment. This may reflect genuine shared involvement of geniculate and pulvinar projections to V5, residual tract overlap despite conservative segmentation, or limited statistical power.

These findings are consistent with the idea that residual geniculo-extrastriate pathways contribute to visual recovery after striate cortex damage. They also align with previous work showing microstructural change in the LGN-V5 pathway following visual training, despite differences in training paradigms (Willis et al., 2024). While both training protocols shared the use of moving targets in the blind visual field, there were also several differences. The present study used forced-choice discrimination of temporally modulated spatial gratings, with luminance contrast of targets dropping as performance increased. In contrast, previous training studies have used moving dots in the blind visual field in a principal direction discrimination task, with task difficulty adapted by reducing motion coherence. The convergence of findings across different training methods suggests that LGN-V5 pathway microstructure may be relevant to visual plasticity more generally, rather than being specific to a single behavioural task. Although the precise contribution of neighbouring tracts remains difficult to disentangle, the recurrence of training-related change in a comparable V5-directed pathway location supports the idea that visual rehabilitation may engage residual geniculo-extrastriate circuitry even in chronic hemianopia.

## Brain-behaviour relationships

Brain-behaviour analyses provide preliminary evidence that LGN-V5 microstructure may be behaviourally meaningful. Baseline FA in LGN-V5 showed the strongest associations with visual outcome, particularly post-training Gabor detection. FA change in LGN-V5 also showed the strongest relationship with improvement in Gabor detection, although this did not survive correction for multiple comparisons. These results should therefore be interpreted cautiously, but are consistent with the possibility that preserved geniculo-extrastriate microstructure may support responsiveness to visual training.

The relationship with clinical perimetry was less clear. We did not identify a robust diffusion MRI marker that predicted change in perimetric sensitivity. This was consistent with our previous fMRI findings (Ajina et al., 2021) and with an independent study investigating neural markers of dot discrimination training (Willis et al., 2024). This may partly reflect differences between outcome measures. The Gabor task used a forced-choice design and may be sensitive to residual visual processing in the absence of full conscious perception, whereas Humphrey perimetry depends on detecting and reporting a brief static light stimulus. These measures may therefore capture partly distinct aspects of visual recovery. In the present study, FA change in LGN-V1 showed the strongest relationship with change in perimetry, raising the possibility that improvements in conscious light detection may depend on different mechanisms from improvements in forced-choice Gabor detection. However, given the small sample size, this interpretation remains speculative.

## Subcortical pathways involving the superior colliculus

A secondary aim of the study was to examine whether purely subcortical pathways between the superior colliculus and LGN or pulvinar showed evidence of training-related change. These pathways are of interest because residual vision and blindsight have been linked to multiple subcortical routes, with evidence supporting roles for LGN (Schmid et al., 2010) and for pulvinar and superior colliculus (Kinoshita et al., 2019; Rodman et al., 1989) with recent work lending support to a multi-pathway model (Takakuwa et al., 2021). Animal studies have also suggested that the superior colliculus may provide visual input to koniocellular neurons in LGN, offering a possible route by which collicular signals could influence geniculo-extrastriate processing (Harting et al., 1991; Lachica and Casagrande, 1993).

In the present study, we did not find significant FA changes in either SC-LGN or SC-pulvinar pathways. However, this negative result should be interpreted with caution. These tracts are short and closely adjacent, with our results demonstrating that they share substantial spatial overlap. This leaves limited portions from which pathway-specific metrics can be extracted. In addition, diffusion tractography cannot establish synaptic directionality or distinguish direct from polysynaptic anatomical routes. The absence of a significant FA change therefore does not exclude a role for superior colliculus in residual vision. Rather, it suggests that the current diffusion MRI approach may be better suited to detecting changes in longer subcortico-cortical pathways rather than in very short, overlapping subcortical connections.

## Limitations and future directions

The main limitation of this study is the small sample size. This restricted statistical power, particularly for brain-behaviour correlations and for analyses involving multiple pathways and outcome measures. For this reason, the present findings are viewed as preliminary and hypothesis-generating. The small cohort also motivated our focus on a single microstructural measure, fractional anisotropy (FA). This metric was preferred over mean diffusivity as it is shown to be sensitive to both degeneration and neuroplasticity (Ramu et al., 2008; Yogarajah et al., 2010). Additional tensor-based metrics such as axial and radial diffusivity could have offered additional prognostic value, but high collinearity renders the need for a much larger sample size. We were also restricted from adopting more complex metrics such as NODDI and mean kurtosis (Fieremans et al., 2011; Zhang et al., 2012) by the limited number of b values in the current dMRI dataset. Such methods could be considered in future, larger studies, as well as possible application of fixel based analyses to consider differences in fibre density (Raffelt et al., 2017).

A second limitation is that tractography of small visual pathways remains technically challenging. Even with conservative overlap-based segmentation, reconstructed pathways may not correspond precisely to discrete anatomical tracts. This is particularly relevant for LGN-V5 and pulvinar-V5 projections, which converge towards a shared cortical target and may interact with wider visual and attentional networks. Nevertheless, by explicitly quantifying overlap and restricting analyses to less ambiguous tract segments, the present study provides a more anatomically cautious framework for studying pathway-specific microstructural change in hemianopia.

In conclusion, this study suggests that visual training in people with hemianopia is associated with increased FA in the LGN-V5 pathway of the lesioned hemisphere, and that microstructure within this pathway may relate to behavioural visual outcome. These findings support a role for geniculo-extrastriate circuitry in visual recovery after stroke, while highlighting that pathway-specific interpretations require caution. More broadly, the study demonstrates the importance of quantifying tract overlap when using diffusion MRI to study small, neighbouring visual pathways. Future work that links multimodal longitudinal imaging with psychophysical and clinical outcomes may help develop methods to stratify patients and tailor visual rehabilitation after stroke.

## Data availability

All anonymised data are available on request from the authors following publication.

## Author Contributions

SA conceived the study, collected the original behavioural and neuroimaging data, supervised the analyses, and contributed to additional analyses. MB carried out the main preliminary analyses under SA’s supervision. AS developed the visual training stimulus and advised on the experimental protocol. HB supervised SA during the original study and contributed to study oversight. RMT contributed to interpretation of the findings and manuscript writing and editing. SA and MB drafted the manuscript. All authors critically reviewed the manuscript and approved the final version.

## Acknowledgements

We thank all the participants for taking part in this study. We also thank the clinical staff who supported with patient recruitment, and the scanning team who assisted in MRI data collection.

---

**Supplementary Figure 1.**
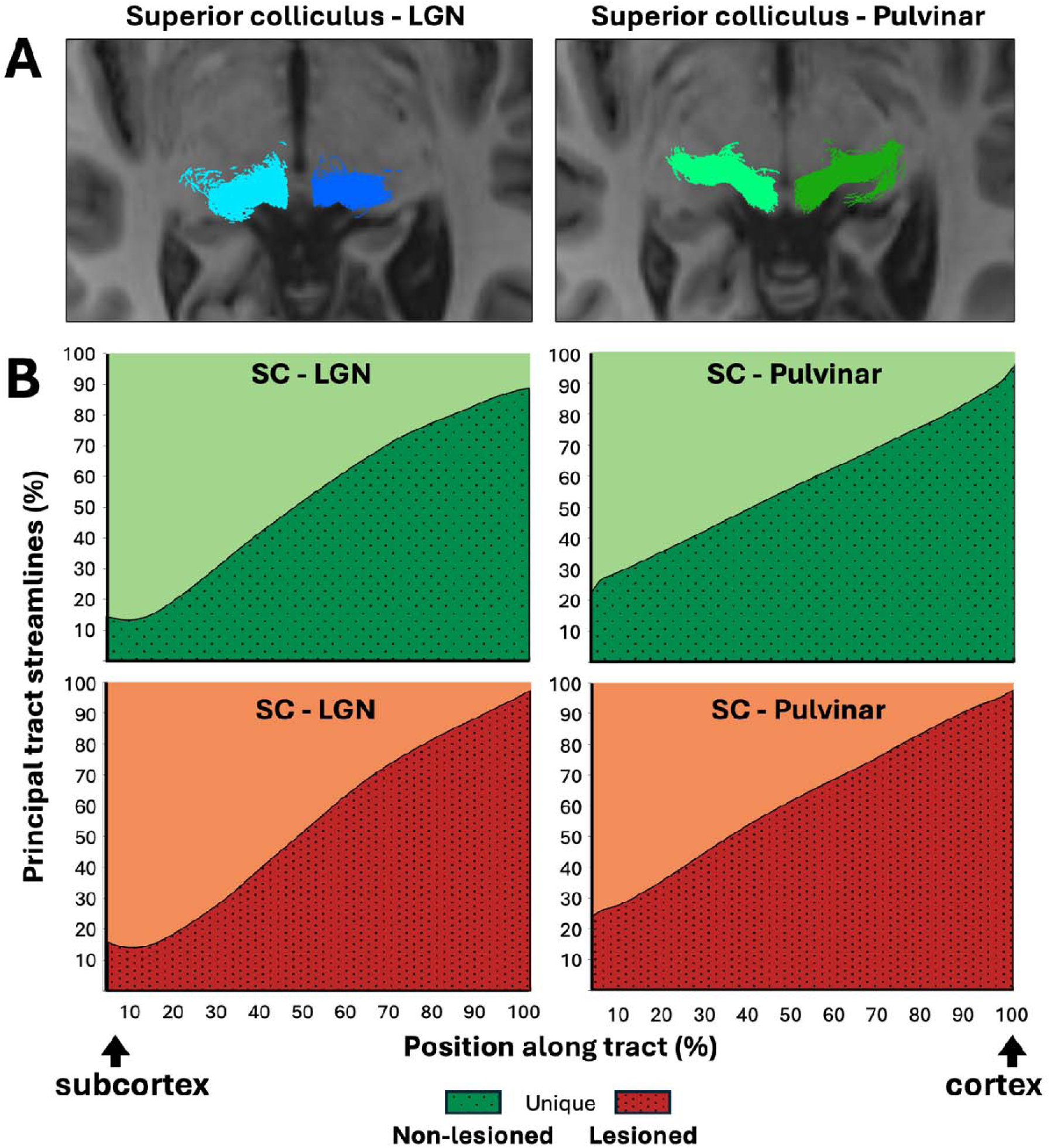
Overlap between superior colliculus (SC)-LGN and SC-pulvinar pathways. Representative reconstructions of the SC-LGN and SC-pulvinar pathways are shown at the top. For each pathway separately, plots show the proportion of streamlines at each node along the tract that were unique to the principal tract or spatially overlapped with the neighbouring subcortical pathway. Position along each tract was normalised from 0 to 100%, with results shown separately for the lesioned and non-lesioned hemispheres. Based on these overlap profiles, nodes 70-100 were selected as the relatively unique portions of the SC-LGN and SC-pulvinar pathways for subsequent microstructural analyses.

**Supplementary Figure 2.**
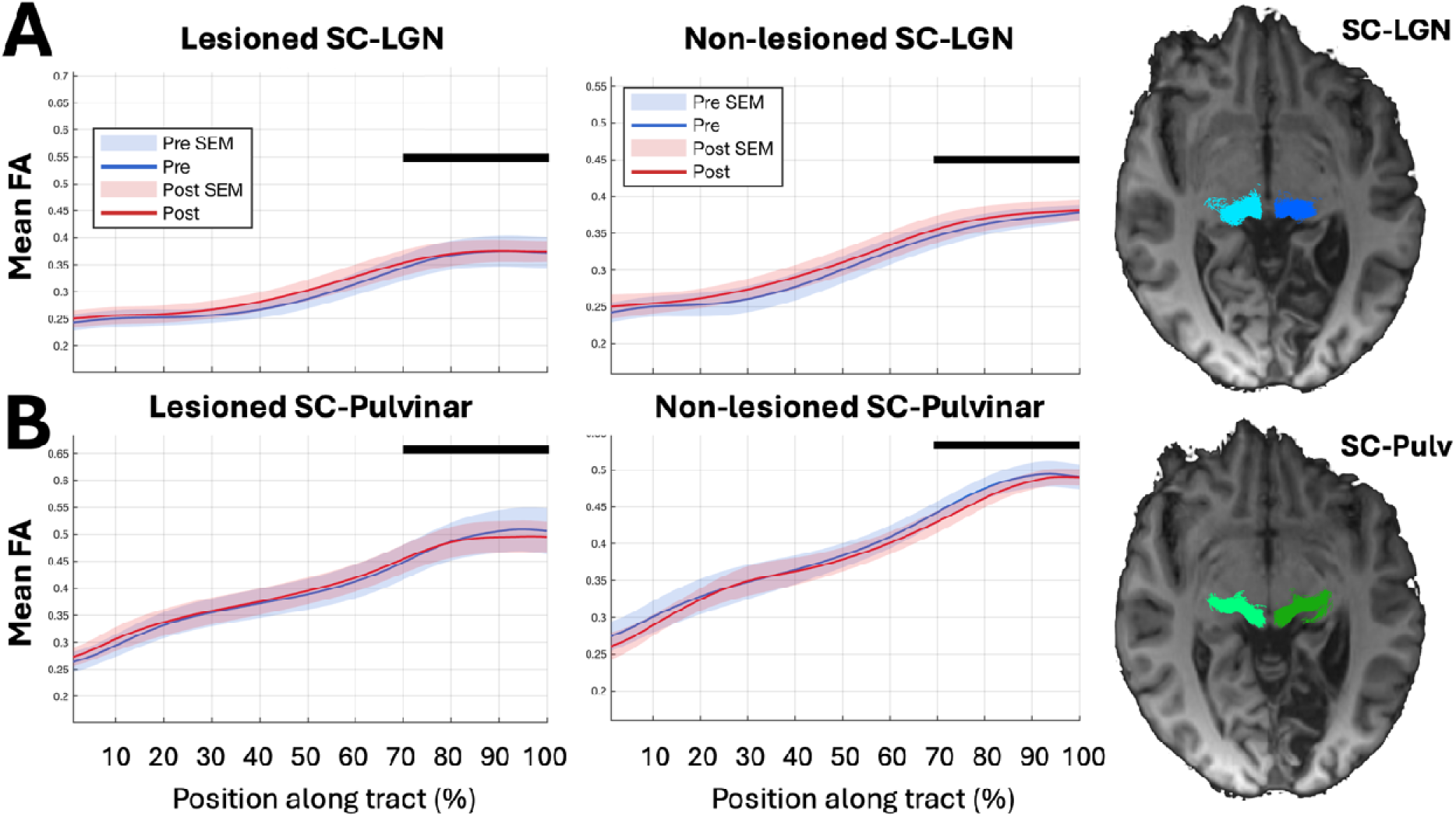
**Fractional anisotropy profiles of superior colliculus (SC)-LGN and SC-pulvinar pathways before and after visual training**. Mean fractional anisotropy (FA) is shown at each position along the **(A)** SC-LGN and **(B)** SC-pulvinar pathways in the lesioned and non-lesioned hemispheres. Position along each tract was normalised from 0 to 100%. Solid lines indicate group mean FA before (blue) and after (red) visual training, with shaded regions representing the standard error of the mean (SEM). Black horizontal bars indicate the tract segments selected for statistical analysis based on the overlap criteria shown in Supplementary Figure 1. Representative reconstructions of each pathway are shown on the right.

## REFERENCES

1. Ajina, S., Bridge, H., 2018. Blindsight relies on a functional connection between hMT+ and the lateral geniculate nucleus, not the pulvinar. PLoS Biology 16. 10.1371/journal.pbio.2005769

2. Ajina, S., Briggs, Farran, Schneider, K., Arcaro, M., Liu, X., Bhatla, N., Marcello, R., n.d. Hidden vision and beyond: Multiscale foundations of thalamic organization. Journal of Neuroscience.

3. Ajina, S., Jünemann, K., Sahraie, A., Bridge, H., 2021. Increased Visual Sensitivity and Occipital Activity in Patients With Hemianopia Following Vision Rehabilitation. J Neurosci 41, 5994–6005. 10.1523/JNEUROSCI.2790-20.2021

4. Ajina, S., Pestilli, F., Rokem, A., Kennard, C., Bridge, H., 2015. Human blindsight is mediated by an intact geniculo-extrastriate pathway. eLife 4. 10.7554/eLife.08935

5. Allen, B., Spiegel, D.P., Thompson, B., Pestilli, F., Rokers, B., 2015. Altered white matter in early visual pathways of humans with amblyopia. Vision Research 114, 48–55. 10.1016/j.visres.2014.12.021

6. Barbot, A., Das, A., Melnick, M.D., Cavanaugh, M.R., Merriam, E.P., Heeger, D.J., Huxlin, K.R., 2021. Spared perilesional V1 activity underlies training-induced recovery of luminance detection sensitivity in cortically-blind patients. Nat Commun 12, 6102. 10.1038/s41467-021-26345-1

7. Basser, P.J., Mattiello, J., Lebihan, D., 1994. Estimation of the Effective Self-Diffusion Tensor from the NMR Spin Echo. Journal of Magnetic Resonance, Series B 103, 247–254. 10.1006/jmrb.1994.1037

8. Beaulieu, C., Allen, P.S., 1994. Determinants of anisotropic water diffusion in nerves. Magnetic Resonance in Med 31, 394–400. 10.1002/mrm.1910310408

9. Beaulieu, C., Does, M.D., Snyder, R.E., Allen, P.S., 1996. Changes in water diffusion due to Wallerian degeneration in peripheral nerve. Magnetic Resonance in Med 36, 627–631. 10.1002/mrm.1910360419

10. Beh, A., McGraw, P.V., Webb, B.S., Schluppeck, D., 2022. Linking Multi-Modal MRI to Clinical Measures of Visual Field Loss After Stroke. Frontiers in Neuroscience 15. 10.3389/fnins.2021.737215

11. Berman, R.A., Wurtz, R.H., 2010. Functional Identification of a Pulvinar Path from Superior Colliculus to Cortical Area MT. J. Neurosci. 30, 6342–6354. 10.1523/JNEUROSCI.6176-09.2010

12. Berman, R.A., Wurtz, R.H., 2008. Exploring the pulvinar path to visual cortex, in: Progress in Brain Research. Elsevier, pp. 467–473. 10.1016/S0079-6123(08)00668-7

13. Bridge, H., Hicks, S.L., Xie, J., Okell, T.W., Mannan, S., Alexander, I., Cowey, A., Kennard, C., 2010. Visual activation of extra-striate cortex in the absence of V1 activation. Neuropsychologia 48, 4148–4154. 10.1016/j.neuropsychologia.2010.10.022

14. Catani, M., Dell’acqua, F., Bizzi, A., Forkel, S.J., Williams, S.C., Simmons, A., Murphy, D.G., Thiebaut de Schotten, M., 2012. Beyond cortical localization in clinico- anatomical correlation. Cortex 48, 1262–1287. 10.1016/j.cortex.2012.07.001

15. Cavanaugh, M.R., Fahrenthold, B.K., Huxlin, K.R., 2025. What V1 Damage Can Teach Us About Visual Perception and Learning. Annu Rev Vis Sci 11, 217–241. 10.1146/annurev-vision-110323-112823

16. Celeghin, A., de Gelder, B., Tamietto, M., 2015. From affective blindsight to emotional consciousness. Consciousness and Cognition 36, 414–425. 10.1016/j.concog.2015.05.007

17. Cowey, A., 2010. The blindsight saga. Experimental Brain Research 200, 3–24. 10.1007/s00221-009-1914-2

18. Das, A., Tadin, D., Huxlin, K.R., 2014. Beyond blindsight: properties of visual relearning in cortically blind fields. J Neurosci 34, 11652–11664. 10.1523/JNEUROSCI.1076-14.2014

19. de Gelder, B., Vroomen, J., Pourtois, G., Weiskrantz, L., 1999. Non-conscious recognition of affect in the absence of striate cortex. Neuroreport 10, 3759–3763. 10.1097/00001756-199912160-00007

20. Dhollander, T., Connelly, A., 2016. A novel iterative approach to reap the benefits of multi-tissue CSD from just single-shell (+b=0) diffusion MRI data.

21. Dhollander, T., Raffelt, D., Connelly, A., 2016. Unsupervised 3-tissue response function estimation from single-shell or multi-shell diffusion MR data without a co-registered T1 image. Presented at the ISMRM Workshop on Breaking the Barriers of Diffusion MRI, p. 5.

22. Fieremans, E., Jensen, J.H., Helpern, J.A., 2011. White matter characterization with diffusional kurtosis imaging. Neuroimage 58, 177–188. 10.1016/j.neuroimage.2011.06.006

23. Glasser, M.F., Coalson, T.S., Robinson, E.C., Hacker, C.D., Harwell, J., Yacoub, E., Ugurbil, K., Andersson, J., Beckmann, C.F., Jenkinson, M., Smith, S.M., Van Essen, D.C., 2016. A multi-modal parcellation of human cerebral cortex. Nature 536, 171–178. 10.1038/nature18933

24. Harting, J.K., Huerta, M.F., Hashikawa, T., van Lieshout, D.P., 1991. Projection of the mammalian superior colliculus upon the dorsal lateral geniculate nucleus: organization of tectogeniculate pathways in nineteen species. J Comp Neurol 304, 275–306. 10.1002/cne.903040210

25. Henriques, R.N., Henson, R., Cam-CAN, Correia, M.M., 2023. Unique information from common diffusion MRI models about white-matter differences across the human adult lifespan. Imaging Neurosci (Camb) 1, imag–1–00051. 10.1162/imag_a_00051

26. Iglesias, J.E., Insausti, R., Lerma-Usabiaga, G., Bocchetta, M., Van Leemput, K., Greve, D.N., van der Kouwe, A., Alzheimer’s Disease Neuroimaging Initiative, Fischl, B., Caballero-Gaudes, C., Paz-Alonso, P.M., 2018. A probabilistic atlas of the human thalamic nuclei combining ex vivo MRI and histology. Neuroimage 183, 314–326. 10.1016/j.neuroimage.2018.08.012

27. Johansen-Berg, H., 2010. Behavioural relevance of variation in white matter microstructure. Current Opinion in Neurology 23, 351–358. 10.1097/WCO.0b013e32833b7631

28. Jones, D.K., Knösche, T.R., Turner, R., 2013. White matter integrity, fiber count, and other fallacies: the do’s and don’ts of diffusion MRI. Neuroimage 73, 239–254. 10.1016/j.neuroimage.2012.06.081

29. Kinoshita, M., Kato, R., Isa, K., Kobayashi, Kenta, Kobayashi, Kazuto, Onoe, H., Isa, T., 2019. Dissecting the circuit for blindsight to reveal the critical role of pulvinar and superior colliculus. Nat Commun 10, 135. 10.1038/s41467-018-08058-0

30. Lachica, E.A., Casagrande, V.A., 1993. The morphology of collicular and retinal axons ending on small relay (W-like) cells of the primate lateral geniculate nucleus. Vis Neurosci 10, 403–418. 10.1017/s0952523800004648

31. Leh, S.E., 2006. Unconscious vision: new insights into the neuronal correlate of blindsight using diffusion tractography. Brain 129, 1822–1832. 10.1093/brain/awl111

32. Leh, S.E., Mullen, K.T., Ptito, A., 2006. Absence of S-cone input in human blindsight following hemispherectomy. European Journal of Neuroscience 24, 2954–2960. 10.1111/j.1460-9568.2006.05178.x

33. Maunsell, J.H., van Essen, D.C., 1983. The connections of the middle temporal visual area (MT) and their relationship to a cortical hierarchy in the macaque monkey. J Neurosci 3, 2563–2586. 10.1523/JNEUROSCI.03-12-02563.1983

34. Pestilli, F., Yeatman, J.D., Rokem, A., Kay, K.N., Wandell, B.A., 2014. Evaluation and statistical inference for human connectomes. Nat Methods 11, 1058–1063. 10.1038/nmeth.3098

35. Pierpaoli, C., Basser, P.J., 1996. Toward a quantitative assessment of diffusion anisotropy. Magn Reson Med 36, 893–906. 10.1002/mrm.1910360612

36. Pollock, A., Hazelton, C., Rowe, F.J., Jonuscheit, S., Kernohan, A., Angilley, J., Henderson, C.A., Langhorne, P., Campbell, P., 2019. Interventions for visual field defects in people with stroke. Cochrane Database of Systematic Reviews. 10.1002/14651858.CD008388.pub3

37. Prabhakar, A.T., Margabandhu, K., Bosco, C.J., Jepegnanam, R.T., Prasad, T., Sampathkumar, S., Sunderraj, E.S., Prasad, J.D., Abraham Ninan, G., Bal, D., Vanjare, H., Jasper, A., Mannam, P., M McKendrick, A., Carter, O., Garrido, M.I., 2026. Pulvinar–posterior superior temporal sulcus connectivity contributes to non-conscious emotion processing in affective blindsight. Cerebral Cortex 36, bhag032. 10.1093/cercor/bhag032

38. Raffelt, D.A., Smith, R.E., Ridgway, G.R., Tournier, J.-D., Vaughan, D.N., Rose, S., Henderson, R., Connelly, A., 2015. Connectivity-based fixel enhancement: Whole-brain statistical analysis of diffusion MRI measures in the presence of crossing fibres. Neuroimage 117, 40–55. 10.1016/j.neuroimage.2015.05.039

39. Raffelt, D.A., Tournier, J.-D., Smith, R.E., Vaughan, D.N., Jackson, G., Ridgway, G.R., Connelly, A., 2017. Investigating white matter fibre density and morphology using fixel-based analysis. Neuroimage 144, 58–73. 10.1016/j.neuroimage.2016.09.029

40. Ramu, J., Herrera, J., Grill, R., Bockhorst, T., Narayana, P., 2008. Brain fiber tract plasticity in experimental spinal cord injury: diffusion tensor imaging. Exp Neurol 212, 100–107. 10.1016/j.expneurol.2008.03.018

41. Riddoch, G., 1917. Dissociation of visual perceptions due to occipital injuries, with especial reference to appreciation of movement. Brain 40, 15–57. 10.1093/brain/40.1.15

42. Rodman, H.R., Gross, C.G., Albright, T.D., 1989. Afferent basis of visual response properties in area MT of the macaque. I. Effects of striate cortex removal. J Neurosci 9, 2033–2050. 10.1523/JNEUROSCI.09-06-02033.1989

43. Rokem, A., Yeatman, J.D., Pestilli, F., Kay, K.N., Mezer, A., Van Der Walt, S., Wandell, B.A., 2015. Evaluating the Accuracy of Diffusion MRI Models in White Matter. PLoS ONE 10, e0123272. 10.1371/journal.pone.0123272

44. Rowe, E.G., Zhang, Y., Garrido, M.I., 2023. Evidence for adaptive myelination of subcortical shortcuts for visual motion perception in healthy adults. Human Brain Mapping 44, 5641–5654. 10.1002/hbm.26467

45. Rowe, F., Walker, M., Rockliffe, J., Pollock, A., Noonan, C., Howard, C., Glendinning, R., Currie, J., 2013. Care provision and unmet need for post stroke visual impairment. The Stroke Association.

46. Rowe, F.J., Hepworth, L.R., Howard, C., Hanna, K.L., Currie, J., 2022. Impact of visual impairment following stroke (IVIS study): a prospective clinical profile of central and peripheral visual deficits, eye movement abnormalities and visual perceptual deficits. Disabil Rehabil 44, 3139–3153. 10.1080/09638288.2020.1859631

47. Sahraie, A., MacLeod, M.J., Trevethan, C.T., Robson, S.E., Olson, J.A., Callaghan, P., Yip, B., 2010. Improved detection following Neuro-Eye Therapy in patients with post-geniculate brain damage. Experimental Brain Research 206, 25–34. 10.1007/s00221-010-2395-z

48. Sahraie, A., Trevethan, C.T., Macleod, M.-J., Weiskrantz, L., Hunt, A.R., 2013. The continuum of detection and awareness of visual stimuli within the blindfield: from blindsight to the sighted-sight. Invest Ophthalmol Vis Sci 54, 3579–3585. 10.1167/iovs.12-11231

49. Sampaio-Baptista, C., Khrapitchev, A.A., Foxley, S., Schlagheck, T., Scholz, J., Jbabdi, S., DeLuca, G.C., Miller, K.L., Taylor, A., Thomas, N., Kleim, J., Sibson, N.R., Bannerman, D., Johansen-Berg, H., 2013. Motor skill learning induces changes in white matter microstructure and myelination. J Neurosci 33, 19499–19503. 10.1523/JNEUROSCI.3048-13.2013

50. Sangchooli, A., Rowe, E.G., Smith, R.E., Dumontheil, roise, Garrido, M.I., 2026. Weakening of subcortical and strengthening of cortical visual pathways across early adolescence. Human Brain Mapping.

51. Schall, J.D., Morel, A., King, D.J., Bullier, J., 1995. Topography of visual cortex connections with frontal eye field in macaque: convergence and segregation of processing streams. J Neurosci 15, 4464–4487. 10.1523/JNEUROSCI.15-06-04464.1995

52. Schmid, M.C., Mrowka, S.W., Turchi, J., Saunders, R.C., Wilke, M., Peters, A.J., Ye, F.Q., Leopold, D.A., 2010. Blindsight depends on the lateral geniculate nucleus. Nature 466, 373–377. 10.1038/nature09179

53. Sen, P.N., Basser, P.J., 2005. A model for diffusion in white matter in the brain. Biophys J 89, 2927–2938. 10.1529/biophysj.105.063016

54. Smith, R.E., Tournier, J.-D., Calamante, F., Connelly, A., 2015. SIFT2: Enabling dense quantitative assessment of brain white matter connectivity using streamlines tractography. Neuroimage 119, 338–351. 10.1016/j.neuroimage.2015.06.092

55. Smith, R.E., Tournier, J.-D., Calamante, F., Connelly, A., 2013. SIFT: Spherical-deconvolution informed filtering of tractograms. Neuroimage 67, 298–312. 10.1016/j.neuroimage.2012.11.049

56. Smith, R.E., Tournier, J.-D., Calamante, F., Connelly, A., 2012. Anatomically-constrained tractography: improved diffusion MRI streamlines tractography through effective use of anatomical information. Neuroimage 62, 1924–1938. 10.1016/j.neuroimage.2012.06.005

57. Sungkarat, W., Chaeyklinthes, T., Thadanipon, K., Plant, G.T., Jindahra, P., 2024. Preserved motion perception and the density of cortical projections to V5 in homonymous hemianopia. Brain Commun 6, fcae436. 10.1093/braincomms/fcae436

58. Takakuwa, N., Isa, K., Onoe, H., Takahashi, J., Isa, T., 2021. Contribution of the Pulvinar and Lateral Geniculate Nucleus to the Control of Visually Guided Saccades in Blindsight Monkeys. J Neurosci 41, 1755–1768. 10.1523/JNEUROSCI.2293-20.2020

59. Tamietto, M., Pullens, P., de Gelder, B., Weiskrantz, L., Goebel, R., 2012. Subcortical Connections to Human Amygdala and Changes following Destruction of the Visual Cortex. Current Biology 22, 1449–1455. 10.1016/j.cub.2012.06.006

60. Torrealba, F., Partlow, G.D., Guillery, R.W., 1981. Organization of the projection from the superior colliculus to the dorsal lateral geniculate nucleus of the cat. Neuroscience 6, 1341–1360. 10.1016/0306-4522(81)90192-5

61. Tournier, J.-D., Calamante, F., Connelly, A., 2010. Improved probabilistic streamlines tractography by 2nd order integration over fibre orientation distributions. Proc. Intl. Soc. Mag. Reson. Med. (ISMRM) 18.

62. Tournier, J.-D., Smith, R., Raffelt, D., Tabbara, R., Dhollander, T., Pietsch, M., Christiaens, D., Jeurissen, B., Yeh, C.-H., Connelly, A., 2019. MRtrix3: A fast, flexible and open software framework for medical image processing and visualisation. Neuroimage 202, 116137. 10.1016/j.neuroimage.2019.116137

63. Warner, C.E., Goldshmit, Y., Bourne, J.A., 2010. Retinal afferents synapse with relay cells targeting the middle temporal area in the pulvinar and lateral geniculate nuclei. Frontiers in Neuroanatomy. 10.3389/neuro.05.008.2010

64. Warner, C.E., Kwan, W.C., Bourne, J.A., 2012. The early maturation of visual cortical area MT is dependent on input from the retinorecipient medial portion of the inferior pulvinar. Journal of Neuroscience 32, 17073–17085. 10.1523/JNEUROSCI.3269-12.2012

65. Weiskrantz, L., Warrington, E.K., Sanders, M.D., Marshall, J., 1974. Visual capacity in the hemianopic field following a restricted occipital ablation. Brain 97, 709–728. 10.1093/brain/97.1.709

66. Willis, H.E., Caron, B., Cavanaugh, M.R., Starling, L., Ajina, S., Pestilli, F., Tamietto, M., Huxlin, K.R., Watkins, K.E., Bridge, H., 2024. Rehabilitating homonymous visual field deficits: white matter markers of recovery-stage 2 registered report. Brain Commun 6, fcae323. 10.1093/braincomms/fcae323

67. Willis, H.E., Cavanaugh, M.R., 2023. Recommended Changes to Standard of Care for Monitoring of Cortically Blind Fields. Policy Insights from the Behavioral and Brain Sciences 10, 308–316. 10.1177/23727322231196563

68. Willis, H.E., Hameed, J., Starling, L., Kaltenbach, C., Moore, M.J., Khan, A., Maxwell, R., Tamietto, M., Ajina, S., Bridge, H., 2025. Voxel-Based Lesion-Symptom Mapping Localizes Residual Visual Function in Hemianopia. J Neurosci 45, e1263242024. 10.1523/JNEUROSCI.1263-24.2024

69. Yeatman, J.D., Dougherty, R.F., Myall, N.J., Wandell, B.A., Feldman, H.M., 2012. Tract Profiles of White Matter Properties: Automating Fiber-Tract Quantification. PLoS ONE 7, e49790. 10.1371/journal.pone.0049790

70. Yogarajah, M., Focke, N.K., Bonelli, S.B., Thompson, P., Vollmar, C., McEvoy, A.W., Alexander, D.C., Symms, M.R., Koepp, M.J., Duncan, J.S., 2010. The structural plasticity of white matter networks following anterior temporal lobe resection. Brain 133, 2348–2364. 10.1093/brain/awq175

71. Zhang, H., Schneider, T., Wheeler-Kingshott, C.A., Alexander, D.C., 2012. NODDI: practical in vivo neurite orientation dispersion and density imaging of the human brain. Neuroimage 61, 1000–1016. 10.1016/j.neuroimage.2012.03.072

